# Drug susceptibility testing and mortality in patients treated for tuberculosis in high-burden countries

**DOI:** 10.1101/370056

**Authors:** Kathrin Zürcher, Marie Ballif, Lukas Fenner, Sonia Borrell, Peter M. Keller, Joachim Gnokoro, Olivier Marcy, Marcel Yotebieng, Lameck Diero, E. Jane Carter, Neesha Rockwood, Robert J. Wilkinson, Helen Cox, Nicholas Ezati, Alash’le G. Abimiku, Jimena Collantes, Anchalee Avihingsanon, Kamon Kawkitinarong, Miriam Reinhard, Rico Hömke, Robin Huebner, Sebastien Gagneux, Erik C. Böttger, Matthias Egger, on behalf of the International Epidemiology Databases to Evaluate AIDS (IeDEA)

**Author notes:** **Correspondence:** Professor Matthias Egger, Institute of Social and Preventive Medicine (ISPM), University of Bern, Mittelstrasse 43, CH-3012 Bern, Switzerland. Equal contribution (Kathrin Zürcher, Marie Ballif, Sebastien Gagneux, Erik C. Böttger, Matthias Egger). **Sources of support** This research was supported by the Swiss National Science Foundation (grant numbers 153442, 310030_166687 and 174281), the National Institutes of Allergy and Infectious Diseases (NIAID) under award numbers U01 AI096299, U01 AI069919, U01 AI069924, U01 AI069911, U01 AI069907, U01 AI096186, and U01 AI069923, and Swiss National Center for Mycobacteria, University of Zurich, Switzerland.

## Abstract

**Background:** Drug resistance and HIV co-infection are challenges for the global control of tuberculosis.

**Methods:** We collected *Mycobacterium tuberculosis* isolates from adult patients in Côte d’Ivoire, Democratic Republic of the Congo, Kenya, Nigeria, South Africa, Peru, and Thailand, stratified by HIV status and tuberculosis drug resistance. Molecular or phenotypic drug susceptibility testing (DST) was done locally and at the Swiss tuberculosis reference laboratory. We examined mortality during treatment according to DST results and treatment adequacy in logistic regression models adjusting for sex, age, sputum microscopy and HIV status.

**Findings:** 634 tuberculosis patients were included; median age was 33.2 years, 239 (37.7%) were female, 272 (42.9%) HIV-positive and 69 (10.9%) patients died. Based on the reference laboratory DST, 394 (62.2%) strains were pan-susceptible, 45 (7.1%) mono-resistant, 163 (25.7%) multidrug-resistant (MDR-TB), and 30 (4.7%) had pre-extensive or extensive drug resistance (pre-XDR/XDR-TB). Results of reference and local laboratories were discordant in 121 (19.1%) cases, corresponding to a sensitivity of 84.3% and a specificity of 90.8%. In patients with drug-resistant tuberculosis, discordant results were associated with increased mortality (risk ratio 1.81; 95% CI 1.07-3.07). In logistic regression, compared to adequately treated patients with pan-susceptible strains, the adjusted odds ratio for death was 4.23 (95% CI 2.16-8.29) for adequately treated patients with drug-resistant strains and 21.54 (95% CI 3.36-138.1) for inadequately treated patients with drug-resistant strains. HIV status was not associated with mortality.

**Interpretation:** Using a reference laboratory standard, inaccurate DST leading to inappropriate treatment of drug-resistant tuberculosis, but not HIV infection, contributed to mortality.

## RESEARCH IN CONTEXT

### Evidence before this study

Multidrug-resistant tuberculosis (MDR-TB) and extensively drug-resistant tuberculosis (XDR-TB) are serious threats to the World Health Organization’s End-TB strategy, due to limited access to rapid drug resistance identification and appropriate treatment for patients with MDR-TB or XDR-TB in many high tuberculosis burden countries. We searched PubMed for systematic reviews and original research articles published in any language up to March 31, 2018. We combined terms for “tuberculosis”, “drug resistance testing”, and “mortality”. Several individual studies and systematic reviews have documented the poor outcomes of MDR-TB and pre-XDR/XDR-TB in high-burden countries. Two Cochrane reviews evaluated the accuracy of molecular tests detecting specific mutations associated with resistance, for example the Xpert MTB/RIF, which is recommended by the World Health Organization to detect rifampicin resistance directly from sputum.

### Added value of this study

To our knowledge, this is the first multi-country study assessing the accuracy of drug susceptibility testing (DST) in routine settings in high-burden countries by comparing local DST results with those from a tuberculosis reference laboratory, and assessing the impact on mortality. The study showed that the accuracy of local DST in high-burden countries was moderate (sensitivity 84%, specificity 91%). Results from the reference and local laboratories were discordant in about 20% of patients. Mortality during treatment was increased almost two-fold in patients with discordant DST results compared to patients with concordant results. Mortality ranged from 6.0% in adequately treated patients with pan-susceptible strains to 53.3% in inadequately treated patients with drug-resistant strains. In multivariable analyses, associations with mortality changed little after adjustment for sex, age, sputum microscopy result and HIV status. Of note, HIV infection was not associated with mortality during tuberculosis treatment.

### Implications of all the available evidence

Drug-resistant tuberculosis is difficult to diagnose and to treat, particularly in high-burden settings, where resources are limited. In these settings, inaccurate DST leading to inappropriate treatment contributes to the high mortality associated with drug-resistant tuberculosis. Access to detailed DST of first- and second-line drugs is required to improve outcomes in patients with MDR-TB and pre-XDR/XDR-TB. Whole genome sequencing is the most promising approach to reach this goal, but much work remains to be done to make this approach feasible and affordable in high-burden countries.

## INTRODUCTION

Tuberculosis is a global public health concern. In 2016, an estimated 10.4 million individuals developed active tuberculosis worldwide, of whom an estimated 1.0 million (10%) were HIV-positive [1]. The scale-up of antiretroviral combination therapy (ART) has substantially improved the prognosis of HIV-positive patients [2,3], and reduced the incidence of tuberculosis in this population [4,5]. However, the risk of tuberculosis among HIV-positive patients on ART remains four times higher than among HIV-negative patients [6].

The emergence of multidrug-resistant tuberculosis (MDR-TB) and extensively drug-resistant tuberculosis (XDR-TB) is another threat to the control of tuberculosis [7– 9]. In 2016, it was estimated that 4% of the new patients and 19% (up to 48% in Eastern Europe) of previously treated patients had MDR-TB [1]. Treatment of MDR-TB and XDR-TB is challenging due to the longer treatment duration, adverse effects and lower efficacy of second-line drugs [10,11]. Strategies to prevent drug-resistant tuberculosis include surveillance, drug susceptibility testing (DST) and ensuring rapid initiation and completion of full courses of effective treatment regimens [12,13]. Culture-based phenotypic DST is considered the gold-standard, but is time and resource intensive, and too slow to influence decisions on starting treatment [14]. Molecular-based resistance testing offers an alternative to culture-based DST [15]. Xpert MTB/RIF (Cepheid, Sunnyvale, CA, USA) detects resistance to rifampicin directly from sputum and provides results within 1.5 hours [16], while line-probe assays (LPAs) from sputum detect resistance to isoniazid, rifampicin, ethambutol, fluoroquinolones, or second-line injectable drugs (aminoglycosides and capreomycin) and provide results within 1-2 days [15].

We compared the results of resistance testing performed locally in ART and tuberculosis programmes in high tuberculosis burden countries to those from gold standard phenotypic DST performed in the Swiss reference laboratory, and examined mortality in HIV-positive and HIV-negative tuberculosis patients with concordant and discordant test results.

## METHODS

This study was part of a larger research project on the evolution of drug-resistant *Mycobacterium tuberculosis* in the context of HIV co-infection within the International Epidemiology Databases to Evaluate AIDS (IeDEA), a global network of ART programs (see www.iedea.org) [17, 18]. Isolates and clinical data were collected from tuberculosis patients in seven high-burden countries in sub-Saharan Africa, Asia and Latin America.

### Patient recruitment and data collection

We included adult patients aged 16 years or older who were treated for active pulmonary tuberculosis in Côte d’Ivoire, Democratic Republic of the Congo (DRC), Kenya, Nigeria, South Africa, Peru, and Thailand. All seven countries are defined by the World Health Organization (WHO) as high tuberculosis burden countries, and DRC, Kenya, Nigeria South Africa and Thailand are also high MDR-TB burden and high HIV/tuberculosis burden countries [19].

HIV-positive tuberculosis patients were recruited from ART clinics participating in IeDEA, HIV-negative patients from tuberculosis clinics serving the same population. Clinics were asked to contribute pulmonary *Mycobacterium tuberculosis* isolates from 25 or more patients within each of the four strata defined by HIV status (positive or negative) and drug resistance (MDR or pan-susceptible). Supplemental <u>Table S1</u> summarizes the characteristics of participating sites. Clinical data were collected online in French or English using the Research Electronic Data Capture (REDCap) tool [20], including age, sex, country, HIV status, CD4 cell count at start of tuberculosis treatment (if HIV positive), sputum smear microscopy result, risk factors for tuberculosis, type of TB patient as defined by WHO, treatment regimen and outcomes.

### Outcomes

Treatment outcomes were defined according to WHO as cured, treatment completed, treatment failure, death, lost to follow-up, transferred to other clinics, ongoing treatment at the time of evaluation or unknown treatment outcome [21]. “Treatment success” included cured patients and patients who completed treatment [21]. The main outcome for this study was mortality during tuberculosis treatment. Outcome data received up to March 31, 2018 were included in analyses.

### Drug susceptibility testing

DST was performed locally using liquid or solid cultures or molecular methods: Xpert MTB/RIF or LPAs, such as Genotype MTBDR*plus* or MTBDR*s*/ tests (Hain Lifesciences, Germany). The reference laboratory of the Swiss National Center for Mycobacteria, Zurich, Switzerland performed DST using the Mycobacteria Growth Indicator Tube liquid medium system (MGIT, Becton Dickinson, USA) with the following drug concentrations: 0.1 mg/L for isoniazid, 1.0 mg/L for rifampicin, 100.0 mg/L for pyrazinamide, 5.0 mg/L for ethambutol, 1.0 mg/L for amikacin and 0.25 mg/L for moxifloxacin, in line with the critical concentrations recently published by WHO [22].

WHO defines mono-resistance as resistance to one first-line anti-tuberculosis drug (isoniazid, rifampicin, pyrazinamide, or ethambutol); MDR as resistance to isoniazid and rifampicin; pre-XDR as MDR with additional resistance to any fluoroquinolone or one of the second-line injectable drugs (amikacin, capreomycin, or kanamycin); XDR as MDR with additional resistances to any fluoroquinolone and at least one of the second-line injectable drugs [21]. The category “other” drug resistance included any other combination. We defined “pan-susceptible” tuberculosis as no resistance against the six drugs tested at the reference laboratory and any resistance as resistance against at least one of the tested drugs. First-line regimens (standard treatment) included first-line anti-tuberculosis drugs (isoniazid, rifampicin, pyrazinamide, and ethambutol) and second-line regimens included a combination of first-line and second-line drugs [21,23].

### Exposure definition and data analysis

We calculated test accuracy statistics for the diagnosis of any drug resistance. We further classified comparisons between the phenotypic and molecular DST results obtained in the local laboratories and the reference laboratory as follow: concordant results, discordance potentially leading to under treatment, discordance potentially leading to over treatment, and other discordant results. We defined drug regimens received by patients as adequate or inadequate based on the reference DST results, taking WHO and local guidelines into account [21]. For example, adequate treatment included first-line regimens for pan-susceptible or mono-resistant tuberculosis other than rifampicin mono-resistance. Second line-regimens prescribed to rifampicin mono-resistant patients, MDR-TB and pre-XDR/XDR-TB patients were classified as adequately treated according to the reference DST results. Inadequate treatment included first-line regimens given to rifampicin mono-resistant patients, MDR-TB and pre-XDR/XDR-TB patients, or second-line regimens given to pan-susceptible tuberculosis patients and mono-resistant patients other than rifampicin mono-resistance [21]. Supplemental <u>Table S2</u> shows the classification of regimen adequacy.

We used descriptive statistics to describe patient characteristics by HIV status and levels of drug resistance based on DST performed at the reference laboratory. We examined determinants of mortality using logistic regression. Patients with unknown or missing treatment outcome, ongoing treatment, missing treatment regimen, missing sputum microscopy and “other” drug-resistant tuberculosis were excluded from regression analyses. Logistic models were adjusted for age, sex, sputum microscopy result and HIV status, and allowed for within-country correlation of standard errors.

Other variables, for example smoking history, diabetes, substance abuse and contact to other tuberculosis patients worsened the fit of the model. For HIV-positive individuals, models were additionally adjusted for CD4 cell count at tuberculosis treatment start. All analyses were done using STATA version 15 (Stata Corporation, College Station, Texas, USA).

### Ethical statement

Local institutional review boards or ethics committees approved the study at all participating sites. Informed consent was obtained where requested per local regulations. The study was also approved by the Cantonal Ethics Committee in Bern, Switzerland.

## RESULTS

We obtained *Mycobacterium tuberculosis* isolates from 871 patients diagnosed between 2013 and 2016. We excluded 237 patients from analyses of the accuracy of DST, mainly because isolates were contaminated or not viable, and a further 61 patients from analyses of mortality, mainly because treatment was ongoing or outcomes unknown at the time of closing the database (<u>Figure 1</u>).

**Figure 1:**
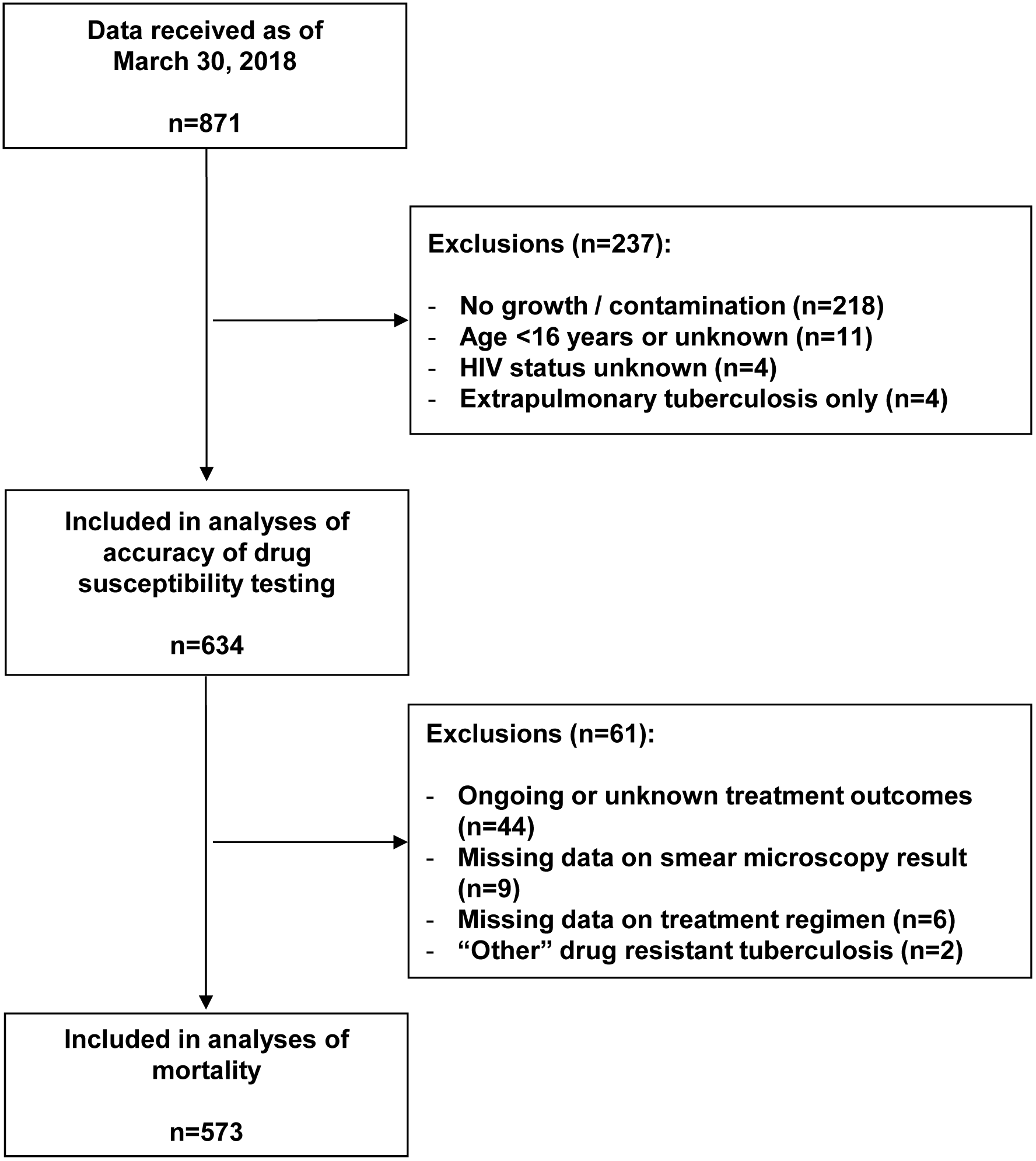
Selection of the study population.

### Characteristics of patients and isolates

The median age of study participants was 33.2 years (interquartile range [IQR] 26.9-42.5 years); 239 (37.7%) were female. The reference laboratory identified 394 (62.1%) pan-susceptible *Mycobacterium tuberculosis* strains, 45 (7.1%) mono-resistant strains, 163 (25.7%) MDR strains, 30 (4.7%) pre-XDR/XDR strains, and 2 (0.3%) strains with other drug resistance profiles (<u>Table 1</u>). Among the 163 patients with MDR-TB, 85 (52.1%) had resistance to rifampicin and isoniazid only, while the remaining patients were additionally resistant to pyrazinamide and/or ethambutol. Among the 24 patients with pre-XDR-TB, resistance to moxifloxacin (n=15) was more frequent than resistance to amikacin (n=9; <u>Table 3</u>). Patients with resistant strains were more likely to receive second-line tuberculosis treatment, and to experience unfavourable treatment outcomes than patients with pan-susceptible strains (<u>Table 1</u>).

**Table 1:** Patient characteristics by phenotypic drug resistance profiles obtained at the Swiss National Center for Mycobacteria.

|  | Pan-susceptible | Any resistance | P-value | Mono-resistance |  |  | Poly-resistance |  |  |
| --- | --- | --- | --- | --- | --- | --- | --- | --- | --- |
|  |  |  |  | INH | RIF | PZA | MDR | Pre-XDR/XDR | Other |
| **Total** | 394 (100) | 240 (100) |  | 29 (100) | 14 (100) | 2 (100) | 163 (100) | 30 (100) | 2 (100) |
| **Sex** |  |  |  |  |  |  |  |  |  |
| Female | 150 (38.1) | 89 (37.1) | 0.80 | 6 (20.7) | 3 (21.4) | 0 | 65 (39.9) | 14 (46.7) | 1 (50.0) |
| Male | 244 (61.9) | 151 (62.9) |  | 23 (79.3) | 11 (78.6) | 2 (100) | 98 (60.1) | 16 (53.3) | 1 (50.0) |
| **Age** | 34.6<br>(27.8-44.6) | 31.5<br>(25.3-40.2) | 0.003 | 34.3<br>(26.5-43.2) | 27.1<br>(24.9-35.5) | 26.1<br>(23.3-28.9) | 31.5<br>(25.4-41.4) | 30.3<br>(24.2-37.5) | 27.3<br>(24.4-30.2) |
| **HIV status** |  |  |  |  |  |  |  |  |  |
| Negative | 200 (50.8) | 162 (67.5) | <0.001 | 20 (69.0) | 8 (57.1) | 1 (50.0) | 114 (69.9) | 18 (60.0) | 1 (50.0) |
| Positive | 194 (49.2) | 78 (32.5) |  | 9 (31.0) | 6 (42.9) | 1 (50.0) | 49 (30.1) | 12 (40.0) | 1 (50.0) |
| **CD4 count at baseline, median (IQR), cells/μl** | 215<br>(85-369) | 161<br>(61-369) | 0.79 | 92.5<br>(55-161) | 63.5<br>(43-81) | 43 | 259<br>(151-528) | 32<br>(5-105) | 213 |
| **No. of observations (%)** | 155 (39.3) | 45 (18.9) |  | 6 (20.7) | 6 (42.9) | 1 (50.0) | 24 (14.7) | 7 (23.3) | 1 (50.0) |
| **Treatment regimen** |  |  |  |  |  |  |  |  |  |
| First line | 369 (93.7) | 46 (19.2) | <0.001 | 27 (93.1) | 0 | 2 (5.4) | 14 (9.2) | 2 (6.7) | 1 (50.0) |
| Second line | 25 (6.3) | 188 (78.3) |  | 2 (6.9) | 14 (100) | 0 | 143 (85.3) | 28 (93.3) | 1 (50.0) |
| Unknown | 0 | 6 (2.5) |  | 0 | 0 | 0 | 6 (5.5) | 0 | 0 |
| **Treatment outcomes** |  |  |  |  |  |  |  |  |  |
| Success | 287 (72.8) | 124 (51.7) | <0.001 | 15 (51.7) | 7 (50.0) | 0 | 88 (54.0) | 13 (43.3) | 1 (50.0) |
| Mortality | 24 (6.1) | 45 (18.8) |  | 7 (24.1) | 2 (14.3) | 1 (50.0) | 24 (14.7) | 10 (33.3) | 1 (50.0) |
| Treatment failure | 12 (3.0) | 10 (4.2) |  | 0 | 0 | 1 (50.0) | 5 (3.1) | 4 (13.3) | 0 |
| Lost to follow-up | 29 (7.4) | 30 (12.5) |  | 1 (3.5) | 3 (21.4) | 0 | 26 (16.0) | 0 | 0 |
| Transfer | 15 (3.8) | 14 (5.8) |  | 0 | 2 (14.3) | 0 | 9 (5.5) | 3 (10.0) | 0 |
| Ongoing treatment / unknown | 27 (6.9) | 17 (7.1) |  | 6 (20.7) | 0 | 0 | 11 (6.7) | 0 | 0 |
| **Country** |  |  |  |  |  |  |  |  |  |
| Côte d'Ivoire | 48 (12.2) | 51 (21.3) | <0.001 | 3 (10.3) | 0 | 0 | 44 (27.0) | 4 (13.3) | 0 |
| Democratic Republic of the Congo | 33 (8.4) | 29 (12.1) |  | 0 | 1 (7.1) | 0 | 19 (11.7) | 9 (30.0) | 0 |
| Kenya | 24 (6.1) | 11 (4.6) |  | 2 (6.9) | 1 (7.1) | 0 | 8 (4.9) | 0 | 0 |
| Nigeria | 20 (5.1) | 36 (15.0) |  | 1 (3.5) | 5 (35.7) | 0 | 26 (16.0) | 4 (13.3) | 0 |
| Peru | 66 (16.8) | 38 (15.8) |  | 8 (27.6) | 0 | 0 | 27 (16.6) | 3 (10.0) | 0 |
| South Africa | 130 (33.0) | 57 (23.8) |  | 6 (20.7) | 7 (50.0) | 1 (50.0) | 32 (15.5) | 10 (33.3) | 1 (50.0) |
| Thailand | 73 (18.5) | 18 (7.5) |  | 9 (31.0) | 0 | 1 (50.0) | 7 (4.3) | 0 | 1 (50.0) |

Analysis based on 634 patients. Numbers (%) or medians (interquartile range) are shown.
INH, isoniazid; MDR, multidrug resistant; PZA, pyrazinamide; RIF, rifampicin; XDR, extensively drug resistant.

**Table 2:** Drug resistance profiles identified at the Swiss National Center for Mycobacteria.

| **Resistance profiles** | **No. of patients** |
| --- | --- |
|  | (n=634) |
| Pan-susceptible | 394 (62.2%) |
| Mono-resistance | 45 (7.1%) |
| INH mono-resistance | 29 |
| RIF mono-resistance | 14 |
| PZA mono-resistance | 2 |
| MDR | 163 (25.7%) |
| INH+RIF | 85 |
| INH+RIF+EMB | 11 |
| INH+RIF+PZA | 47 |
| INH+RIF+EMB+PZA | 20 |
| Pre-XDR | 24 (3.2%) |
| INH+RIF +MOX+EMB+PZA | 8 |
| INH+RIF +MOX+EMB | 1 |
| INH+RIF +MOX+PZA | 4 |
| INH+RIF +MOX | 2 |
| INH+RIF +AMK+PZA+EMB | 4 |
| INH+RIF +AMK+PZA | 4 |
| INH+RIF +AMK | 1 |
| XDR | 6 (0.8%) |
| INH+RIF +AMK+MOX+EMB | 3 |
| INH+RIF +AMK+MOX+PZA | 2 |
| INH+RIF +AMK+MOX | 1 |
| Other | 2 (0.3%) |
| INH+MOX | 1 |
| INH+PZA | 1 |

AMK, amikacin; EMB, ethambutol; INH, isoniazid; MDR, multidrug resistant; MOX, moxifloxacin; PZA, pyrazinamide; RIF, rifampicin; XDR, extensively drug resistant.

**Table 3:** Concordance and discordance of drug susceptibility results obtained from reference and local laboratories.

| Concordance/<br>discordance of<br>DST results | DST results by laboratory |  | Total<br>(n=634) | Test used at local laboratories |  |  |  |
| --- | --- | --- | --- | --- | --- | --- | --- |
|  | Reference laboratory<br>(phenotypic) | Local laboratories |  | Xpert<br>MTB/RIF<sup>a</sup> | Culture | LPA | Combination<br>of tests |
| **Concordance** | Pan-susceptible | Pan-susceptible | 332 | 167 | 101 | 1 | 5 |
|  | RIF mono-resistance | RIF mono-resistance | 8 | 0 | 0 | 0 | 7 |
|  | INH mono-resistance | INH mono-resistance | 8 | 0 | 8 | 0 | 0 |
|  | MDR | MDR | 153 | 49 | 44 | 8 | 52 |
|  | Pre-XDR and XDR | Pre-XDR and XDR | 12 | 0 | 1 | 2 | 9 |
|  | Total |  | 513 (80.9) | 216 (42.1) | 154 (30.0) | 11 (2.1) | 73 (14.2) |
| **Discordance**<br>**potentially**<br>**leading to under**<br>**treatment** | MDR | Pan-susceptible | 5 | 2 | 2 | 0 | 1 |
|  | Pre-XDR and XDR | MDR | 18 | 6 | 7 | 0 | 5 |
|  | Total |  | 23 (3.6) | 8 (34.8) | 9 (39.1) | 0 | 6 (26.1) |
| **Discordance**<br>**potentially**<br>**leading to over**<br>**treatment** | Pan-susceptible | RIF mono-resistance | 14 | 0 | 0 | 3 | 9 |
|  | Pan-susceptible | MDR | 14 | 3 | 8 | 0 | 3 |
|  | Pan-susceptible | Other mono-resistance<sup>b</sup> | 33 | 2 | 31 | 0 | 1 |
|  | Other mono-resistance<sup>c</sup> | MDR | 5 | 0 | 5 | 0 | 0 |
|  | MDR | Pre-XDR or XDR | 1 | 0 | 0 | 0 | 1 |
|  | Total |  | 67 (10.6) | 5 (7.5) | 44 (6.6) | 3 (4.5) | 14 (20.9) |
| **Other**<br>**discordance** | Pan-susceptible | EMB, SM | 1 | 0 | 1 | 0 | 0 |
|  | RIF mono-resistance | MDR | 7 | 2 | 0 | 0 | 5 |
|  | Other mono-resistance<sup>d</sup> | Pan-susceptible | 17 | 13 | 3 | 0 | 0 |
|  | INH, MOX | Mono-resistance | 1 | 0 | 1 | 0 | 0 |
|  | IHN, PZA | MDR | 1 | 0 | 1 | 0 | 0 |
|  | MDR | RIF mono-resistance | 3 | 0 | 0 | 1 | 2 |
|  | MDR | EMB, SM | 1 | 1 | 0 | 0 | 0 |
|  | Total |  | 31 (4.9) | 16 (51.6) | 6 (19.4) | 1 (3.2) | 7 (22.6) |

Analysis based on 634 patients. Number of patients (%) are shown.
DST, drug susceptibility testing; EMB, ethambutol; INH, isoniazid; LPA, line probe assay; MDR, multidrug resistance; PZA, pyrazinamide; RIF, rifampicin; SM, streptomycin; XDR, extensively drug resistant.
In a few patients, the test used to diagnose drug-resistant infection at the local laboratories and the treatment regimen was unknown. Therefore, numbers do not always add up to the row totals.
<sup>a</sup> RIF resistance diagnosed with Xpert MTB/RIF only was categorized as MDR, since those patients were treated as MDR-TB patients.
<sup>b</sup> Twenty-one strains were resistant to EMB, ten to SM and two INH.
<sup>c</sup> Five strains were resistant to INH.
<sup>d</sup> Fifteen strains were resistant to INH, two to PZA

A total of 272 (42.9%) tuberculosis patients were HIV-positive, with a median CD4 cell count at the start of tuberculosis treatment of 192 cells/μl (IQR 77.5-369 cells/μl). Among them, 175 (64.3%) were either on ART at the start of tuberculosis treatment or initiated ART within 3 months; the ART status of the remaining patients was unknown. Compared to HIV-negative individuals, HIV-positive patients were more likely to be female, more likely to have both pulmonary and extrapulmonary disease, and more likely to be patients with recurrent tuberculosis (supplemental <u>Table S3</u>). HIV-positive patients were also more likely to have a negative sputum smear microscopy result and more likely to have a pan-susceptible *Mycobacterium tuberculosis* infection than HIV-negative patients.

### Drug susceptibility testing and treatments

Local laboratories used the Xpert MTB/RIF system, LPAs, phenotypic DST or a combination of these methods to diagnose drug-resistant infections and inform treatment regimens (<u>Table 3</u>, supplemental <u>Table S1</u>). Sensitivity and specificity for the detection of any drug resistance were 84.3% and 90.8%, respectively. The likelihood ratio was 9.2 (95% CI 6.2-13.7) for a DST indicating resistance positive test and 0.17 (0.14-0.22) for a negative test; accuracy was 86.8% (83.9-89.3%).

Results from the reference laboratory and local laboratories were concordant for 513 (80.9%) and discordant for 121 (19.1%) patients. There were 23 (3.6%) discrepancies potentially leading to under treatment, 67 (10.6%) discordant results potentially leading to over treatment, and 31 (4.9%) other discordances (<u>Table 3</u>, supplementary <u>Table S2</u>). When analysing the treatments received, they were adequate in 491 of 507 (96.8%) patients with concordant DST results compared to 94 of 121 patients (77.7%) with discordant results (P<0.001).

### Mortality

After excluding 61 (9.6%) patients with unknown treatment outcomes, missing data or “other” drug resistance (<u>Figure 1</u>), mortality ranged from 9.9% among patients with concordant DST results to 40.9% among patients with discordant results potentially leading to under treatment. Mortality was 6.4% in pan-susceptible tuberculosis, 25.6% in mono-resistant tuberculosis, 16.4% in MDR-TB and 34.5% in pre-XDR/XDR-TB cases (<u>Figure 2</u>, <u>Table 3</u>). In patients with pan-susceptible *Mycobacterium tuberculosis* strains, mortality was 5.9% (18/307) if DST results were concordant between the reference laboratory and local laboratories and 10.0% (6/60) if DST results were discordant (P=0.24). In patients with drug-resistant strains, mortality was 17.0% (29/171) if DST results were concordant, but 30.8% (16/52) if DST results were discordant (P=0.030). The risk ratio comparing discordant with concordant results was 1.81 (95% CI 1.07-3.07), and the population attributable fraction 16.0%. Mortality increased from 5.95% (20/336) in adequately treated patients with pan-susceptible tuberculosis to 53.3% (8/15) in patients with drug-resistant strains receiving inadequate treatment (<u>Figure 2</u>, <u>Table 3</u>).

**Figure 2:**
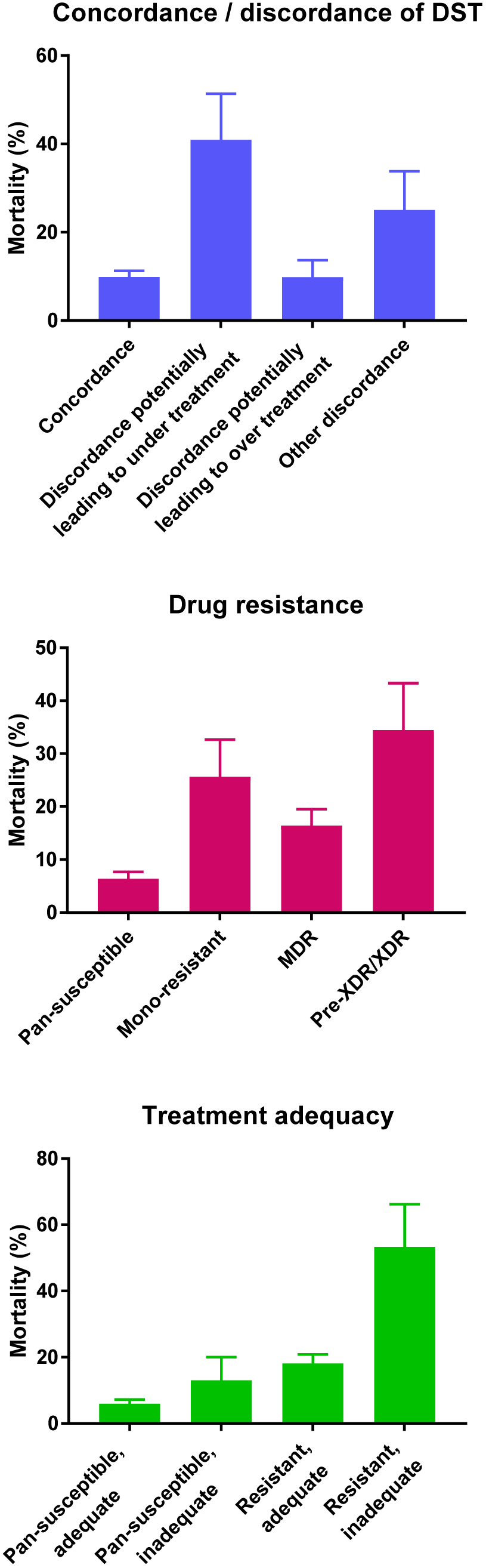
Mortality according to drug resistance, to concordance or discordance of drug susceptibility testing (DST) results and to treatment adequacy. Error bars are standard errors. P-values <0.001 for difference in mortality across categories. Analysis based on 573 patients.

In multivariable logistic models adjusted for sex, age, sputum microscopy result and HIV status, discordant DST results continued to be associated with increased mortality compared to concordant DST results (<u>Table 4</u>). Compared to concordant DST results, the adjusted odds ratio (aOR) of death was 9.53 (95% CI 1.04-87.32) for patients with discordant results potentially leading to under treatment. Similarly, drug resistance was associated with higher mortality compared to pan-susceptible tuberculosis. The aOR was 4.67 (95% Cl 2.59-8.41) for any type of drug resistance, and 11.3 (95% 2.41-53.3) for pre-XDR/XDR (<u>Table 4</u>). Finally, compared to adequately treated patients with pan-susceptible strains, the aOR for death was 4.23 (95% CI 2.16-8.29) for adequately treated patients with resistant strains and 21.54 (95% CI 3.36-138.08) for patients with resistant strains receiving inadequate regimens (<u>Table 4</u>). Sex, positive sputum smear microscopy and HIV status were not associated with the odds of death. The results from univariable models were similar to the aOR from multivariable models (<u>Table S4</u>). When restricting the analysis to HIV-positive patients, mortality was higher among patients with CD4 cell counts <50 cells/μL: the aOR was 3.50 (95% CI 1.27-9.64) compared to patients with higher CD4 counts at tuberculosis treatment start.

**Table 4:** Results from logistic regression models of the probability of death during tuberculosis treatment.

|  | No. of patients | No. of deaths (%) | Model 1<br>aOR (95% CI) | Model 2<br>aOR (95% CI) | Model 3<br>aOR (95% CI) |
| --- | --- | --- | --- | --- | --- |
| **Concordance / discordance of DST results** |  |  |  |  |  |
| Concordance | 466 | 46 (9.9) | 1 |  |  |
| Discordance potentially leading to under treatment | 22 | 9 (40.9) | 9.53 (1.04-87.32) |  |  |
| Discordance potentially leading to over treatment | 61 | 6 (9.8) | 1.01 (0.26-3.88) |  |  |
| Other discordance | 24 | 6 (25.0) | 4.40 (2.14-9.03) |  |  |
| **Drug resistance <sup>a</sup>** |  |  |  |  |  |
| Pan-susceptible | 359 | 23 (6.4) |  | 1 |  |
| Mono-resistance | 39 | 10 (25.6) |  | 5.38 (2.62-11.04) |  |
| MDR | 146 | 24 (16.4) |  | 3.43 (1.91-6.16) |  |
| Pre-XDR/XDR | 29 | 10 (34.5) |  | 11.33 (2.41-53.3) |  |
| **Treatment adequacy by drug resistance** |  |  |  |  |  |
| Pan-susceptible, adequate | 336 | 20 (6.0) |  |  | 1 |
| Pan-susceptible, inadequate | 23 | 3 (13.0) |  |  | 2.81 (0.50-15.81) |
| Any resistance, adequate | 199 | 36 (18.1) |  |  | 4.23 (2.16-8.29) |
| Any resistance, inadequate | 15 | 8 (53.3) |  |  | 21.54 (3.36-138.08) |
| **Sex** |  |  |  |  |  |
| Female | 219 | 20 (9.1) | 1 | 1 | 1 |
| Male | 354 | 47 (13.3) | 1.50 (0.84-2.67) | 1.48 (0.83-2.66) | 1.55 (0.83-2.88) |
| **Age** (per 1 year increase) | 573 | 67 (11.7) | 1.04 (1.02-1.06) | 1.04 (1.03-1.06) | 1.04 (1.02-1.07) |
| **Sputum microscopy** |  |  |  |  |  |
| Negative | 111 | 10 (9.0) | 1 | 1 | 1 |
| Positive | 462 | 57 (12.3) | 1.41 (0.40-5.00) | 1.45 (0.47 -4.44) | 1.25 (0.34-4.58) |
| **HIV status** |  |  |  |  |  |
| Negative | 337 | 43 (12.8) | 1 | 1 | 1 |
| Positive | 236 | 24 (10.2) | 1.04 (0.51-2.09) | 0.85 (0.42-1.69) | 1.07 (0.53-2.16) |

Models based on 573 patients with complete data for all variables shown.
Model 1 was adjusted for concordance / discordance of DST results, sex, age, sputum microscopy and HIV status; model 2 was adjusted for drug resistance, sex, age, sputum microscopy and HIV status; model 3 was adjusted for treatment adequacy, sex, age, sputum microscopy and HIV status.
Abbreviations: DST, drug susceptibility testing; MDR, multidrug resistant; XDR, extensively drug-resistant
<sup>a</sup> Results from the Swiss National Reference Center for Mycobacteria

## DISCUSSION

This study of patients treated for drug-resistant or drug-susceptible tuberculosis in seven high tuberculosis burden countries showed that the accuracy of DST testing in routine care was moderate, with discordant results from local DST compared to phenotypic DST in a reference laboratory in about 20 percent of patients. Discordant results led to inadequate treatment and contributed to the excess mortality associated with drug-resistant tuberculosis. As expected, mortality increased with the degree of drug resistance and was higher in patients who received inadequate treatment regimens. To our knowledge, this is the first study to assess the accuracy of DST in real world, routine settings and to examine the impact of inaccurate results on mortality. Our findings support the recent call for a precision medicine approach to the treatment of drug-resistant tuberculosis, guided by detailed DST, to replace the standardised, empirical combination regimens used in many high tuberculosis burden low- and middle-income countries [24].

At present, WHO recommends that “Xpert MTB/RIF should be used as the initial diagnostic test in individuals suspected of having MDR-TB or HIV-associated tuberculosis” [25], based on a Cochrane review of test accuracy studies in adults with suspected rifampicin-resistance or MDR-TB [26]. In line with this recommendation, Xpert MTB/RIF was the most commonly used test in our study sites. The Cochrane review reported a pooled sensitivity of 95%, based on 17 studies and 555 patients with rifampicin-resistant strains [26]. The pooled specificity was 98%. We examined accuracy of DST strategies at the level of the local laboratories in high-burden countries, in routine care settings, rather than by evaluating a single test. Our estimates of sensitivity and specificity, for the detection of any drug resistance, were considerably lower (84.3% and 90.8%, respectively), despite the fact that, in some patients, a combination of more than one test was used (generally Xpert MTB/RIF followed by LPA or by culture).

There are concerns both about false-negative and false-positive Xpert MTB/RIF test results, and a policy of confirmatory testing has been introduced in South Africa and Brazil [27,28]. The discordant DST results that potentially led to under treatment of drug-resistant tuberculosis (false negative for resistance) were mainly based on locally performed cultures, Xpert MTB/RIF tests, or a combination of the two. Of note, the recently developed Xpert MTB/RIF Ultra assay has been shown to improve detection of rifampicin resistance [29]. Culture-based tests dominated the discordances that potentially led to over treatment, while Xpert MTB/RIF dominated in the category of discordances with unclear clinical significance. We acknowledge that some discordances could be explained by mixed infections, heteroresistance, or minority resistant populations [30,31].

LPAs were rarely used in our study, possibly because they have been widely replaced by Xpert MTB/RIF, which is easier to use and provides results in a shorter time. In addition, LPA suffer from suboptimal accuracy for isoniazid resistance, and WHO recommends that culture-based DST for isoniazid should still be used, particularly in patients with suspected MDR-TB where the LPA result does not detect isoniazid resistance [32]. In one case, the local laboratory detected resistance to ethambutol but this could not be confirmed in the reference laboratory: DST is challenging for ethambutol and less reproducible [33].

Data on treatment outcomes in drug-resistant tuberculosis are scarce, particularly for sub-Saharan Africa. A recent systematic review of treatment outcomes in MDR-TB included data on mortality among adults from seven studies from sub-Saharan Africa, six from South Africa and one from Lesotho [34]. In these studies, mortality during tuberculosis treatment ranged from 12.4% in patients with MDR-TB treated in a referral hospital in the Western Cape, South Africa [35], to 45.8% in a study of XDR-TB patients from three South African provinces [36]. Our results extend these data to other countries in the region, and add further data for Peru and Thailand. Furthermore, our study confirms the poor outcome in patients with INH mono-resistant tuberculosis who are treated with first-line regimens (as recommended by WHO during the study period [37]), in line with a study from Durban, South Africa [38] and a recent systemic review and meta-analysis [39].

In patients co-infected by HIV, the treatment of drug-resistant tuberculosis is challenging for several reasons, including the poorer absorption of drugs [40], the risk of the immune reconstitution inflammatory syndrome (IRIS) [41], or interactions between antiretroviral and second-line tuberculosis drugs [42–44]. In contrast to previous studies from South Africa, which reported higher mortality at end of treatment in HIV-positive patients with MDR-TB compared to HIV-negative MDR-TB patients [35,45], we found no association with HIV infection, although confidence intervals were wide. The median CD4 cell count of HIV-positive patients was considerably higher in our study (192 cells/μL) than in the South African studies [35,45], which may explain the discrepant results. A study from Lesotho [46] also found little evidence for a difference in mortality between HIV-positive patients (median CD4 cell count 185 cells/ μL) and HIV-negative patients. Finally, for patients with XDR-TB, treatment outcomes have been uniformly poor in previous studies, irrespective of HIV status [36].

Our study has several limitations. We sampled eligible patients within strata defined by drug resistance and HIV infection, and therefore could not estimate the incidence or prevalence of drug-resistant tuberculosis in HIV-positive or HIV-negative patients. In previous studies, HIV infection has not been consistently associated with drug resistance [27], but it is clear that in regions with a high-burden of HIV, the majority of patients with MDR-TB will be co-infected with HIV [27]. Although we initially exceeded the planned sample size, about a quarter of patients had to be excluded from analyses of drug susceptibility, mainly due to lack of growth or contamination of cultures, and about a third was excluded from the analysis of mortality outcomes, mainly because vital status was unknown at database closure. The reference laboratory tested resistance against six drugs, and we will have missed resistance against other drugs used, for example rifabutin, kanamycin, ethionamide or levofloxacin. Further, the presence of different subpopulations of *Mycobacterium tuberculosis* in isolates tested at the local sites vs reference laboratory might have introduced variability in phenotypic or molecular DST testing [47].

In conclusion, our study shows that the accuracy of DST testing in routine care in high-burden countries was limited and that inaccurate results led to inadequate treatment and contributed to the excess mortality associated with drug-resistant tuberculosis. Our results support the notion that access to detailed DST of first- and second-line drugs at treatment initiation is required to improve outcomes in patients with MDR-TB and pre-XDR/XDR-TB [27]. Whole genome sequencing is the most promising approach to reach this goal, but much work remains to be done to make this approach feasible and affordable in low- and middle-income countries [27]. In particular, direct testing of sputum samples should become routine to circumvent lengthy mycobacterial cultures [39]. A standardised approach for the interpretation of drug resistance conferring mutations has recently been developed [48]. In the meantime, the capacity for the phenotypic and molecular DST testing recommended by WHO should be increased to ensure the most adequate treatment of drug-resistant tuberculosis in these settings.

## ACKNOWLEDGEMENTS

We thank all sites who participated in this survey and the patients whose data were used in this study. We are grateful to the Tuberculosis Working Group of IeDEA for helpful discussions. We also would like to thank all regional data centers, who contributed to the coordination of the study. RJW is supported by the Francis Crick Institute (10218), which is funded by the Wellcome Trust, Cancer Research UK, and Research Councils UK. He also receives support from the Wellcome Trust (104803, 203135). HC is supported by a Wellcome Trust fellowship and reports grants from UK Medical Research Council and the National Research Foundation of South Africa.

## CONFLICTS OF INTEREST

AA has received honoraria fees from Jensen-Cilag, Gilead and Bristol-Myers Squibb. All other authors have no conflicts of interest to declare.

## Funding

National Institutes of Allergy and Infectious Diseases, Swiss National Science Foundation, Swiss National Center for Mycobacteria.

---

**Table S1:**
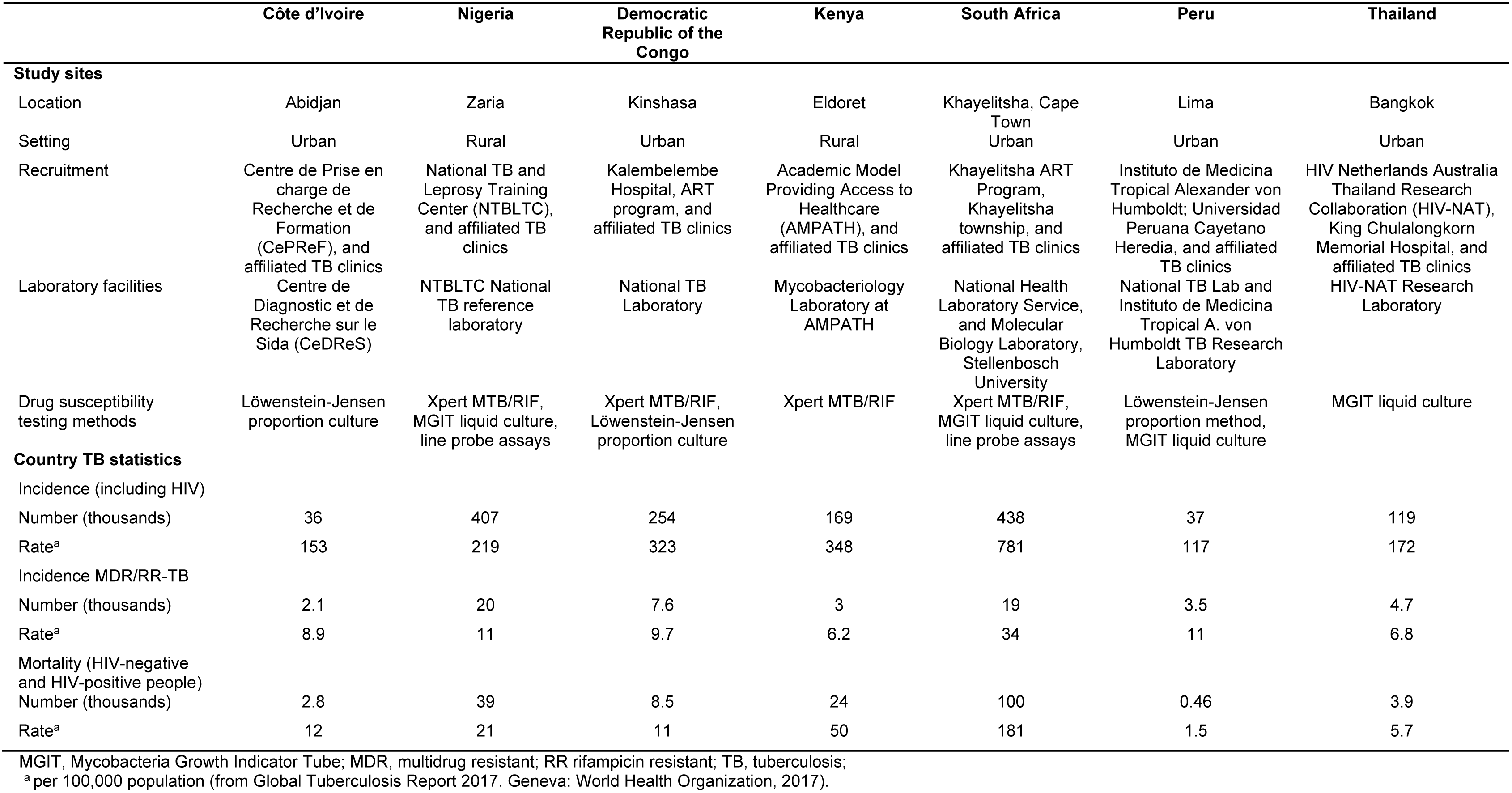
Characteristics of participating study sites and settings.

**Table S2:**
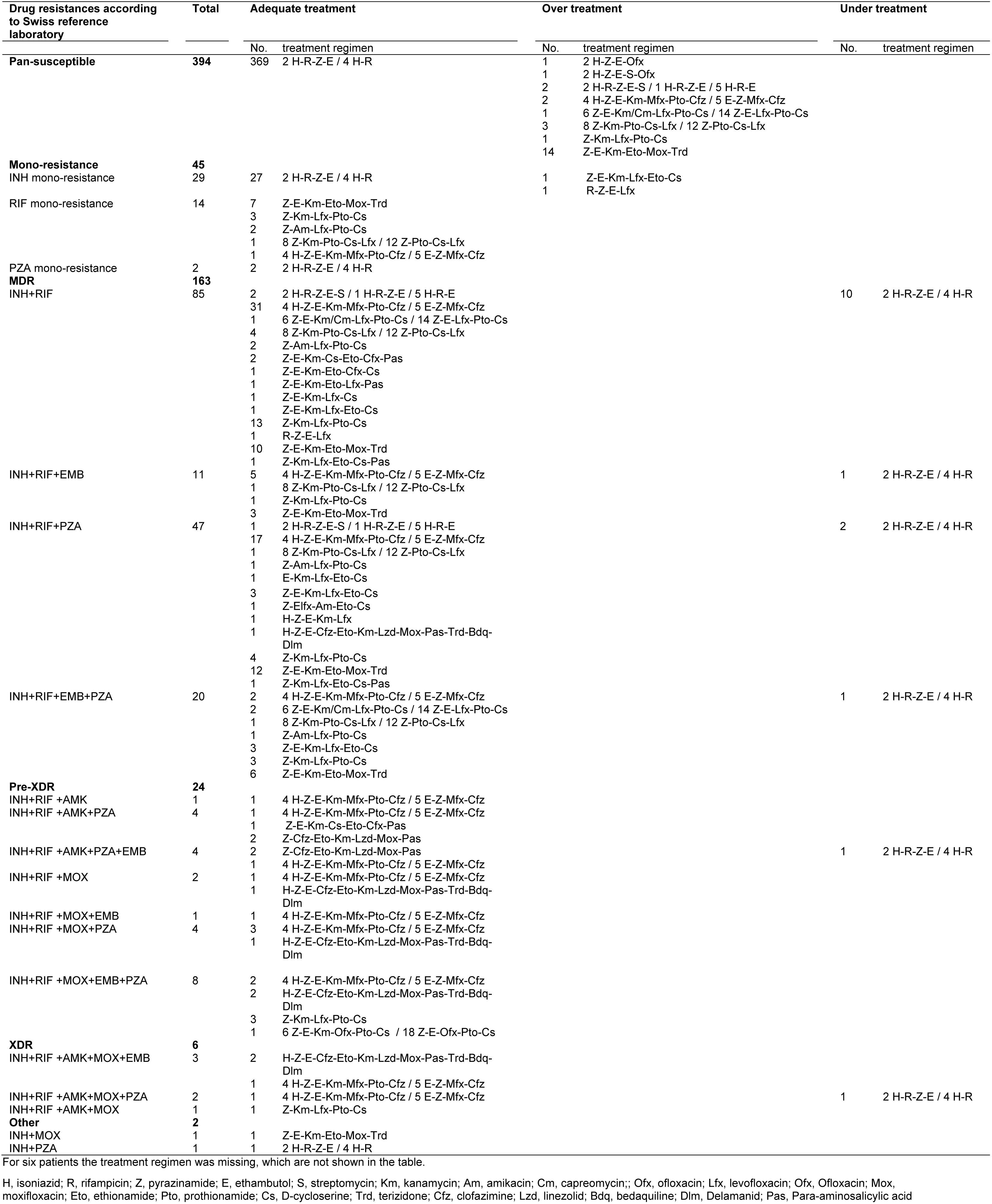
Classification of treatment regimens by drug resistance profile.

**Table S3:** Patient characteristics by HIV status at diagnosis of tuberculosis.

|  | All Patients<br>(n=634) | HIV-negative<br>(n=362) | HIV-positive<br>(n=272) | p-value |
| --- | --- | --- | --- | --- |
| **Age (years)** | 33.2 (26.9-42.5) | 31.7 (25.1-43.3) | 34.7 (29.1-42.0) | 0.49 |
| **Sex** |  |  |  |  |
| Male | 395 (62.3) | 249 (69.8) | 146 (53.7) | <0.001 |
| Female | 239 (37.7) | 113 (31.2) | 126 (46.3) |  |
| **Site of TB disease** |  |  |  |  |
| Pulmonary | 609 (96.1) | 355 (98.1) | 254 (93.4) | 0.003 |
| Pulmonary and extrapulmonary | 25 (3.9) | 7 (1.9) | 18 (6.6) |  |
| **CD4 count at baseline (cells/μl)** | - | - | 192 (77.5-369) |  |
| *No. of observations (%)* | - | - | 200 (73.5) |  |
| **Type of TB patient** |  |  |  | <0.001 |
| New patient | 411 (64.8) | 233 (64.4) | 178 (65.4) |  |
| Recurrent TB | 120 (18.9) | 56 (15.5) | 64 (23.5) |  |
| Treatment after failure | 70 (11.0) | 56 (15.5) | 14 (5.2) |  |
| Treatment after default | 27 (4.3) | 15 (4.1) | 12 (4.4) |  |
| Unknown | 6 (0.9) | 2 (0.5) | 4 (1.5) |  |
| **Sputum smear microscopy** |  |  |  | <0.001 |
| Negative | 113 (17.8) | 46 (12.7) | 67 (24.6) |  |
| Positive | 512 (80.8) | 312 (86.2) | 200 (73.5) |  |
| Unknown | 9 (1.4) | 4 (1.1) | 5 (1.8) |  |
| **TB drug resistance <sup>a</sup>** |  |  |  | <0.001 |
| Pan-susceptible | 394 (62.1) | 200 (55.2) | 194 (71.3) |  |
| Any resistance | 240 (37.9) | 162 (44.8) | 78 (28.7) | <0.001 |
| *Mono-resistant* | 45 (7.1) | 29 (8.0) | 16 (5.9) |  |
| *MDR* | 163 (25.7) | 114 (31.5) | 49 (18.0) |  |
| *Pre-XDR / XDR* | 30 (4.7) | 18 (5.0) | 12 (4.4) |  |
| *Other* | 2 (0.3) | 1 (0.3) | 1 (0.4) |  |
| **TB treatment outcome** |  |  |  | 0.012 |
| Success | 411 (64.8) | 238 (65.7) | 173 (63.6) |  |
| *Cure* | 298 (47.0) | 169 (46.7) | 129 (47.4) |  |
| *Treatment completed* | 113 (17.8) | 69 (19.1) | 44 (16.2) |  |
| Treatment failure | 22 (3.5) | 10 (2.8) | 12 (4.4) |  |
| Death | 69 (10.9) | 43 (11.9) | 26 (9.6) |  |
| Lost to follow-up | 59 (9.3) | 40 (11.0) | 19 (7.0) |  |
| Transfer | 29 (4.6) | 17 (4.7) | 12 (4.4) |  |
| Ongoing treatment | 4 (0.6) | 1 (0.3) | 3 (1.1) |  |
| Unknown | 40 (6.3) | 13 (3.6) | 27 (9.9) |  |
| **Country** |  |  |  | <0.001 |
| Côte d'Ivoire | 99 (15.6) | 57 (15.7) | 42 (15.4) |  |
| Democratic Republic of the Congo | 62 (9.8) | 50 (13.8) | 12 (4.4) |  |
| Kenya | 35 (5.5) | 15 (4.1) | 20 (7.4) |  |
| Nigeria | 56 (8.8) | 37 (10.2) | 19 (7.0) |  |
| Peru | 104 (16.4) | 64 (17.7) | 40 (14.7) |  |
| South Africa | 187 (29.5) | 84 (23.2) | 103 (37.9) |  |
| Thailand | 91 (14.4) | 55 (15.2) | 36 (13.2) |  |

Analysis based on 634 patients. Numbers (%) or medians (interquartile range) are shown. MDR, multidrug resistant; TB, tuberculosis; XDR, extensively drug resistant
<sup>a</sup> Results from the Swiss National Reference Center for Mycobacteria

**Table S4.** Results from univariable logistic regression models of the probability of death during tuberculosis treatment.

|  | No. of patients | No. of deaths (%) | Model 1<br>OR (95% CI) | Model 2<br>OR (95% CI) | Model 3<br>OR (95% CI) |
| --- | --- | --- | --- | --- | --- |
| **Concordance / discordance of DST results** |  |  |  |  |  |
| Concordance | 466 | 46 (9.9) | 1 |  |  |
| Discordance potentially leading to under treatment | 22 | 9 (40.9) | 6.32 (2.56-15.59) |  |  |
| Discordance potentially leading to over treatment | 61 | 6 (9.8) | 1.00 (0.41-2.44) |  |  |
| Other discordance | 24 | 6 (25.0) | 3.04 (1.15-8.05) |  |  |
| **Drug resistance <sup>a</sup>** |  |  |  |  |  |
| Pan-susceptible | 359 | 23 (6.4) |  | 1 |  |
| Mono-resistance | 39 | 10 (25.6) |  | 5.03 (2.19-11.60) |  |
| MDR | 146 | 24 (16.4) |  | 2.87 (1.56-5.28) |  |
| Pre-XDR/XDR | 29 | 10 (34.5) |  | 7.69 (3.21-18.44) |  |
| **Treatment adequacy by drug resistance** |  |  |  |  |  |
| Pan-susceptible, adequate | 336 | 20 (6.0) |  |  | 1 |
| Pan-susceptible, inadequate | 23 | 3 (13.0) |  |  | 2.37 (0.65-8.65) |
| Any resistance, adequate | 199 | 36 (18.1) |  |  | 3.49 (1.96-6.22) |
| Any resistance, inadequate | 15 | 8 (53.3) |  |  | 18.06 (5.95-54.82) |
| **Sex** |  |  |  |  |  |
| Female | 219 | 20 (9.1) | 1 | 1 | 1 |
| Male | 354 | 47 (13.3) | 1.52 (0.88-2.65) | 1.52 (0.88-2.65) | 1.52 (0.88-2.65) |
| **Age** (per 1 year increase) | 573 | 67 (11.7) | 1.03 (1.01-1.05) | 1.03 (1.01-1.05) | 1.03 (1.01-1.05) |
| **Sputum microscopy** |  |  |  |  |  |
| Negative | 111 | 10 (9.0) | 1 | 1 | 1 |
| Positive | 462 | 57 (12.3) | 1.42 (0.70-2.88) | 1.42 (0.70-2.88) | 1.42 (0.70-2.88) |
| **HIV status** |  |  |  |  |  |
| Negative | 337 | 43 (12.8) | 1 | 1 | 1 |
| Positive | 236 | 24 (10.2) | 0.77 (0.46-1.31) | 0.77 (0.46-1.31) | 0.77 (0.46-1.31) |

Models based on 573 patients with complete data for all variables shown.
Abbreviations: DST, drug susceptibility testing; MDR, multidrug resistant; XDR, extensively drug-resistant
<sup>a</sup> Results from the Swiss National Reference Center for Mycobacteria

